# Human MAIT cell cytolytic effector proteins synergize to overcome carbapenem resistance in *Escherichia coli*

**DOI:** 10.1101/2020.01.16.908806

**Authors:** Caroline Boulouis, Wan Rong Sia, Muhammad Yaaseen Gulam, Jocelyn Qi Min Teo, Thanh Kha Phan, Jeffrey Y. W. Mak, David P. Fairlie, Ivan K. H. Poon, Tse Hsien Koh, Peter Bergman, Lin-Fa Wang, Andrea Lay Hoon Kwa, Johan K. Sandberg, Edwin Leeansyah

## Abstract

Mucosa-associated invariant T (MAIT) cells are abundant antimicrobial T cells in humans, and recognize antigens derived from the microbial riboflavin biosynthetic pathway presented by the MHC-Ib-related protein (MR1). However, the mechanisms responsible for MAIT cell antimicrobial activity are not fully understood, and the efficacy of these mechanisms against antibiotic resistant bacteria has not been explored. Here, we show that MAIT cells mediate MR1-restricted antimicrobial activity against *E. coli* clinical strains in a manner dependent on the activity of cytolytic proteins, but independent of production of pro-inflammatory cytokines or induction of apoptosis in infected cells. The combined action of the pore-forming antimicrobial protein granulysin and the serine protease granzyme B released in response to TCR-mediated recognition of MR1-presented antigen is essential to mediate control against both cell-associated and free-living *E. coli*. Furthermore, MAIT cell-mediated bacterial control extend to multidrug-resistant *E. coli* primary clinical isolates additionally resistant to carbapenems, a class of last resort antibiotics. Notably, high levels of granulysin and granzyme B in the MAIT cell secretomes directly damage bacterial cells by increasing their permeability, rendering initially resistant *E. coli* susceptible to the bactericidal activity of carbapenems. These findings define the role of cytolytic effector proteins in MAIT cell-mediated antimicrobial activity, and indicate that granulysin and granzyme B synergize to restore carbapenem bactericidal activity and overcome carbapenem resistance in *E. coli*.

**One Sentence Summary:** Potent antimicrobial activity of human MAIT cells overcomes carbapenem-resistance in control of *Escherichia coli*

## Introduction

Mucosa-associated invariant T (MAIT) cells are innate-like T cells that are highly abundant in mucosal tissues, the liver, lungs and gastrotintestinal tract, and in peripheral blood [1]. MAIT cells are mostly CD8α^+^ [2, 3], express a semi-invariant T cell receptor (TCR), and recognize antigens in complex with the MHC-Ib-related protein (MR1) [4]. MR1 displays an extraordinary level of evolutionary conservation among placental and marsupial mammals [5], strongly supporting the notion that MR1 and MAIT cells perform critical functions in the immune system. The MR1-presented antigens recognized by MAIT cells are derivatives of intermediates in the microbial synthesis of vitamin B_2_ (riboflavin) and are produced by many bacteria [6–8]. Riboflavin is a critical component in a wide variety of bacterial cellular processes [9]. MAIT cells are thus able to recognize and respond to a broad set of bacteria [10]. Following TCR-mediated recognition of MR1-presented bacterial riboflavin metabolite antigens, MAIT cells rapidly mediate a range of effector responses, including cytokine production, cytotoxicity, antimicrobial activity, and tissue repair function [11–18]. The abundance and antimicrobial features of MAIT cells, as well as the highly-conserved nature of MR1, strongly suggest that MAIT cells are important for the protection of the host against bacterial pathogens [19]. Moreover, the conserved nature of the MAIT cell antigens across bacterial species *via* the shared components of the riboflavin biosynthetic pathway and antimicrobial features of MAIT cells, support the hypothesis that they may have the capacity to recognize and respond to drug-resistant bacteria.

Infections caused by antimicrobial-resistant (AMR) bacteria are a serious threat to global public health. In particular, carbapenem-resistant *Enterobacteriaceae* (CRE), including *Escherichia coli, Klebsiella pneumoniae*, and *Enterobacter* spp., have emerged as a serious problem for hospitalized patients [20]. CRE are multidrug-resistant Gram-negative bacteria that have acquired further resistance to one class of the last resort antibiotics, the carbapenems. Resistance to carbapenems in *Enterobacteriaceae* involves multiple mechanisms, including expression of efflux pumps, impermeability due to porin loss, and expression of β-lactamases with the ability to degrade carbapenems [20]. The polymyxins, and more specifically colistin, are the last resort antibiotics currently available to treat CRE infections. However, polymyxin-resistant CRE is on the rise, rendering them extensively drug-resistant (XDR) or even pan-drug resistant (PDR) [21]. The World Health Organization has therefore listed CRE in the critical category requiring further research and development of new treatments [22].

Despite the body of evidence that MAIT cells contribute to bacterial clearance and play a protective role in various bacterial infections [19], the mechanisms of MAIT cell antimicrobial activity remain relatively little explored. Moreover, it is unknown whether MAIT cell antimicrobial activity extends to AMR bacterial pathogens. In this study, we therefore investigated the mechanisms underlying MAIT cell antimicrobial activity by dissecting the roles of cytolytic proteins and cytokines in controlling bacterial growth. We extended these investigations by exploring the ability of MAIT cell cytolytic proteins, released in response to cognate recognition of MR1-presented antigen, to potentiate carbapenem-induced bactericidal activity against primary clinical isolates of *E. coli* from patients with CRE infections. Notably, MAIT cell antimicrobial activity mediated by the combined action of granulysin (Gnly) and Granzyme (Grz) B potently enhanced bactericidal activity of carbapenems against carbapenem-resistant strains of *E. coli.* Our findings thus define the mechanism underlying MAIT cell antimicrobial activity and support the concept that MAIT cell mobilization may protect the host from drug-resistant bacterial pathogens.

## Results

### MAIT cells kill E. coli-infected cells and reduce live bacterial load within infected cells

To study the antimicrobial effector functions of MAIT cells, we developed an approach using the drug-sensitive *E. coli* clinical strain EC120S isolated from a patient suffering from a bloodstream infection [23]. We also developed a method for short-term MAIT cell expansion to provide the cell numbers required to perform such assays (fig. S1A). The *E. coli* strain EC120S infected HeLa cells efficiently as determined by pHrodo red-labelled live bacteria (fig. S1B and C). pHrodo becomes fluorescence in the acidic environment of cellular endosomal compartments, and can therefore be used to evaluate *E. coli* internalization [24]. EC120S infected, replicated, and remained viable inside HeLa cells for at least 24 h post-infection (fig. S1D). MAIT cells rapidly degranulated and killed the tested epithelial cell lines (HeLa and A549) infected with *E. coli* EC120S (Fig. 1A-D and fig. S1E and F) through an MR1-dependent mechanism (Fig. 1C-D).

**Fig. 1.**
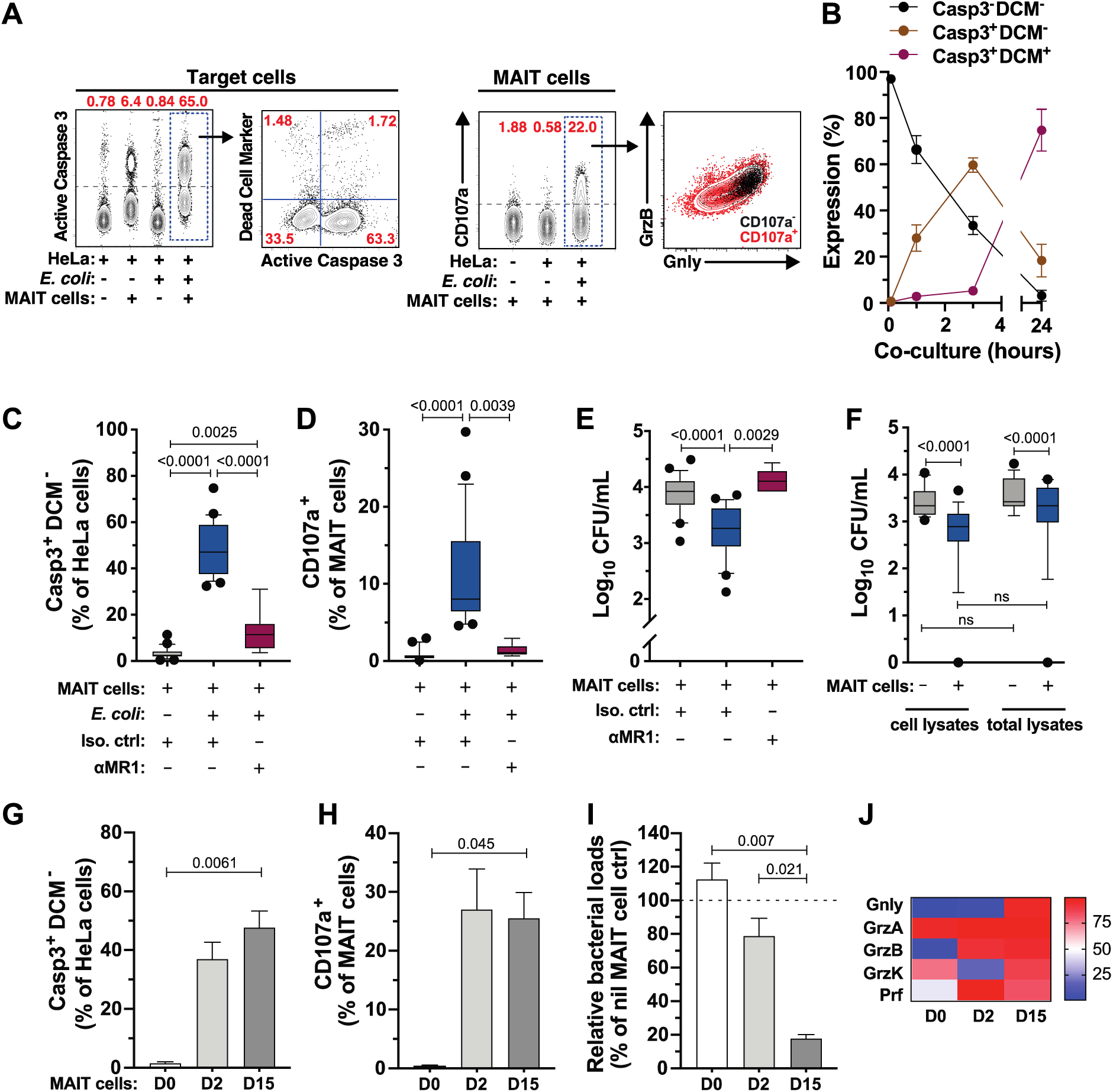
MAIT cells kill bacteria-infected cells and suppress bacterial loads. (A) Assessment of apoptosis of E. coli EC120S-infected HeLa cells by caspase (Casp)3 activity and MAIT cell degranulation by CD107a expression. (B) Apoptosis of infected HeLa cells at indicated time-point (n=11). (C) Measurement of early apoptosis (Casp3^+^DCM-) on uninfected or EC120S-infected HeLa cells, (D) degranulation by MAIT cells, and (E) bacterial counts following lysis of infected HeLa cells after 3 h of co-culture with and without MAIT cells in the presence of anti-MR1 or isotype control (n=8 for EC120S-infected HeLa cells+anti-MR1, n=25 for others). (F) Bacterial counts in infected HeLa cell lysates or in total lysates (cell lysates plus supernatants) after 3 h of co-culture with or without MAIT cells (n=16). (G) Apoptosis of EC120S-infected HeLa cells, (H) degranulation by MAIT cells, and (I) relative bacterial loads following co-culture with MAIT cells (n=4), and (J) percentage of cytolytic proteins expressed by MAIT cells from day (D)0, 2, and 15 of culture (n=3-10). Statistical significance was calculated using mixed-effects analysis followed by Tukey’s multiple comparison test (C-E), Friedman’s test with Dunn’s multiple comparisons test (F), or repeated-measure one-way ANOVA (G-I). ns, not significant.

Because MAIT cells achieved complete lysis (Casp3^+^ DCM^+^) of the infected cells by 24 h, we examined MAIT cell antimicrobial activity after 3 h, when the cell membrane integrity of most infected cells undergoing apoptosis remained intact, as shown by staining with amine-reactive cytoplasmic dyes (Fig. 1B and C). Notably, bacterial viability in both HeLa and A549 cells was significantly reduced in the presence of MAIT cells, *via* an MR1-dependent mechanism (Fig. 1E, fig. S1G). To assess whether the reduced bacterial load reflected true bacterial killing by MAIT cells, or simply the release of bacteria into the supernatants during the assay, the bacterial loads in the cell lysates alone or in total lysates were enumerated. In both conditions, the presence of MAIT cells decreased bacterial load, indicating direct bacterial control (Fig. 1F). Contrary to MAIT cells, Vα7.2^-^ non-MAIT T cells were not able to control bacterial growth (fig. S1H-J). In addition, the capacity of MAIT cells to degranulate and induce apoptosis of infected target cells was superior to that of Vα7.2^-^ T cells (fig. S1H and I). We next investigated whether MAIT cells at resting state were similarly able to mediate antimicrobial activity. Consistent with previous studies [24, 25], resting MAIT cells were inefficient in killing HeLa cells infected with *E. coli* or pulsed with the MR1 ligand 5-(2-oxopropylideneamino)-6-D-ribitylaminouracil (5-OP-RU) (Fig. 1G and H, fig. S2A-C). Interestingly, resting MAIT cells also failed to control bacterial loads in infected cells (Fig. 1I). To determine if the inability of resting MAIT cells to mediate antimicrobial activity was simply due to their weak cytotoxicity, MAIT cells were stimulated with IL-2 + IL-7 for 2 days to allow upregulation of cytotoxic molecules [24, 25]. These cytokine-activated MAIT cells were able to degranulate and kill cells infected with *E. coli* or pulsed with 5-OP-RU, but still failed to efficiently control bacterial loads within infected cells when compared to MAIT cells cultured for 15 days (Fig. 1G-I, fig. S2A-C). Interestingly, this temporal regulation of MAIT cell antimicrobial activity appeared to correspond to the differential expression of cytolytic proteins by MAIT cells (Fig. 1J, fig. S2D and E). These results indicate that antigen-activated MAIT cells have the capacity to kill infected target cells in an MR1-dependent manner through caspase 3 activation, and progressively over several days develop the ability to control bacterial growth within infected cells.

### MAIT cells mediate antimicrobial activity through the cytolytic protein-dependent pathway

Next, the intracellular levels of GrzA, GrzB, GrzK, Gnly, and perforin (Prf) were measured using flow cytometry to determine the transfer of cytolytic effector proteins into *E. coli*-infected cells after co-culture with MAIT cells. With the exception of GrzK, all measured cytolytic proteins were detected in HeLa target cells (Fig. 2A), matching with their expression pattern in MAIT cells (Fig. 1J, fig. S2D and E). The inhibition of bacterial growth associated negatively with cell viability (Fig. 2B), but positively with apoptosis (Fig. 2C) and MAIT cell expression of GrzB (Fig. 2D and E), Gnly (Fig. 2F), and co-expression of GrzB and Gnly (Fig. 2G). Because GrzB was readily detected in *E. coli*-infected target cells co-cultured with MAIT cells (Fig. 2A), we evaluated GrzB activity inside the infected target cells using a fluorescent GrzB substrate (GranToxiLux) (*24*). Active GrzB was detected in HeLa target cells infected with *E. coli* (Fig. 2H), indicating that MAIT cells delivered GrzB into infected cells.

**Fig. 2.**
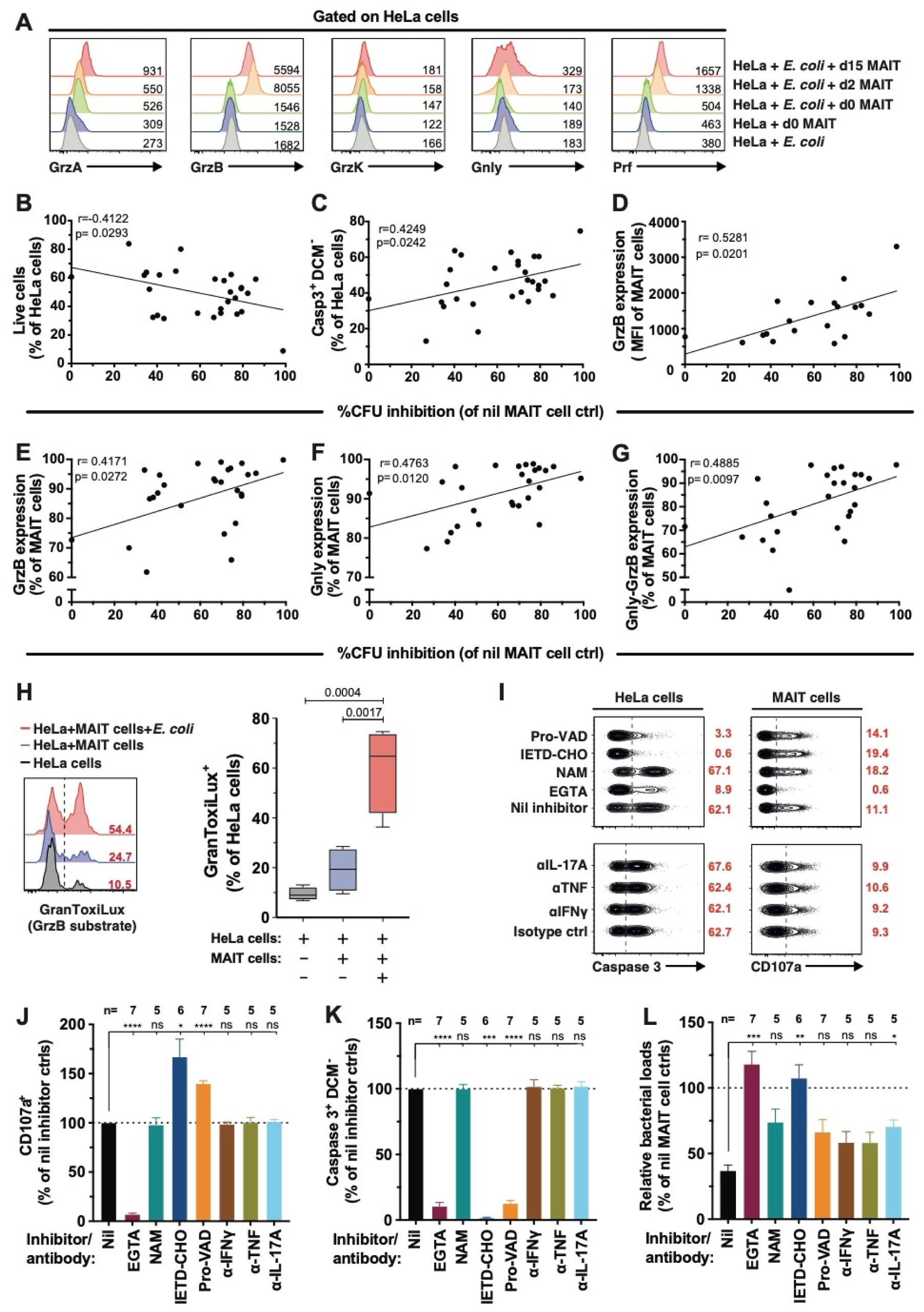
MAIT cell antimicrobial activity is associated with cytolytic protein expression. (A) Detection of MAIT cell-derived cytolytic protein contents in E. coli EC120S-infected or uninfected HeLa cells following 3 h co-culture with MAIT cells obtained at different days following MAIT cell activation. Representative histograms from 4 independent donors are shown. (B) Correlation between live and (C) apoptotic EC120S-infected HeLa cells, (D) GrzB intensity, (E) GrzB expression, (F) Gnly expression, and (G) Gnly-GrzB co-expression in MAIT cells with the inhibition of bacterial growth (n=19 for G, others n=28). (H) Levels of the GrzB substrate GranToxiLux activity in uninfected or EC120S-infected HeLa cells with or without MAIT cells (n=4). (I) Flow cytometry plots of (J) degranulation by MAIT cells, (K) apoptosis in EC120S-infected HeLa cells, and (L) the relative bacterial loads in the presence of the indicated inhibitors or mAbs. Statistical significance was determined using the ordinary one-way ANOVA (H), or mixed-effects analysis (J-L) followed by Dunnett’s or Tukey’s post-hoc test as appropriate. Correlations were calculated using the Spearman test. **** p<0.0001, *** p<0.001, * p<0.05. ns: not significant.

In order to assess MAIT cell use of the cytolytic protein pathway to control bacterial growth, cytolytic protein activity was selectively inhibited by pharmacological inhibitors. EGTA, a Ca^2+^-specific chelator that inhibits the release of cytolytic granules (*25*), was used in the presence of Mg^2+^ supplementation. EGTA strongly decreased MAIT cell degranulation as determined by CD107a expression (Fig. 2I and J), reduced the delivery of cytolytic proteins (fig. S2F) and caspase 3 activation in infected target cells (Fig. 2I and K), and abolished the bacterial control by MAIT cells (Fig. 2L). To investigate the role of GrzA and GrzB in MAIT cell antimicrobial activity, their activation was blocked using nafamostat mesylate (NAM) and Ac-IETD-CHO, respectively [26, 27]. NAM had no effect on MAIT cell degranulation, caspase 3 activation, and bacterial counts (Fig. 2I-L). In contrast, while Ac-IETD-CHO had no effect on MAIT cell degranulation (Fig. 2I and J), it strongly abrogated caspase 3 activation in infected target cells (Fig. 2I and K), and impaired bacterial control (Fig. 2L). To assess whether apoptosis of infected cells also contributed to bacterial control, caspase 3 activation was blocked using the pan-caspase inhibitor Pro-VAD-FMK. The pan-caspase inhibitor had no effect on MAIT cell degranulation (Fig. 2I and J), but severely diminished caspase 3 activation in infected cells without affecting bacterial counts (Fig. 2I, K, L). This suggests that bacterial control by MAIT cells was independent of infected cell apoptosis. Finally, we assessed whether IFNγ, TNF, and IL-17A, pro-inflammatory cytokines produced by MAIT cells (fig. S2G), played a role in MAIT cell bacterial control by blocking with neutralizing Abs. Blocking IFNγ, TNF, or IL-17A did not significantly affect degranulation or caspase 3 activation (Fig. 2I-K). IL-17A blockade, but not IFNγ or TNF blockade, only slightly diminished MAIT cell antimicrobial activity (Fig. 2L). Altogether, these findings indicate that MAIT cells use the cytolytic protein pathway to kill infected cells, and this may play an important role in early control of bacterial loads.

### MAIT cells recognize and mediate antimicrobial activity against carbapenem-resistant E. coli clinical strains

Next, we tested the hypothesis that MAIT cells maintain antimicrobial activity against carbapenem-resistant *E. coli* (CREC) clinical isolates. To address this, we first investigated whether CREC strains EC234, EC241, EC362, and EC385 (Table S1) were riboflavin autotroph by culture in riboflavin assay medium (RAM), a riboflavin-deficient broth. All CREC isolates grew in both RAM and nutrient-rich lysogeny broth at comparable growth rates, and the addition of external riboflavin did not influence their growth (fig. S3A-H). Moreover, the expression of *ribA*, the gene encoding the first enzyme of the riboflavin biosynthesis pathway, was similar among the clinical isolates (fig. S3I). These findings confirmed that the CREC clinical isolates were riboflavin-synthesis competent, a requirement for generating the riboflavin biosynthetically derived MAIT cell antigens [7].

Next, the ability of CREC to stimulate MAIT cells was tested by incubating PBMCs with fixed bacteria for 24 h. All CREC strains induced degranulation and expression of GrzB, IFNγ, TNF, and IL-17A by MAIT cells at comparable levels (Fig. 3A, fig. S3J). Furthermore, MAIT cells displayed similar patterns of polyfunctionality when stimulated with the different CREC strains, although some differences in MAIT cell response profiles were noted (fig. S3K). These variations occurred despite similar uptake of the bacteria by the PBMCs (fig. S3L). MAIT cell responses to CREC were predominantly MR1-dependent although this differed somewhat between the strains (Fig. 3B).

**Fig. 3.**
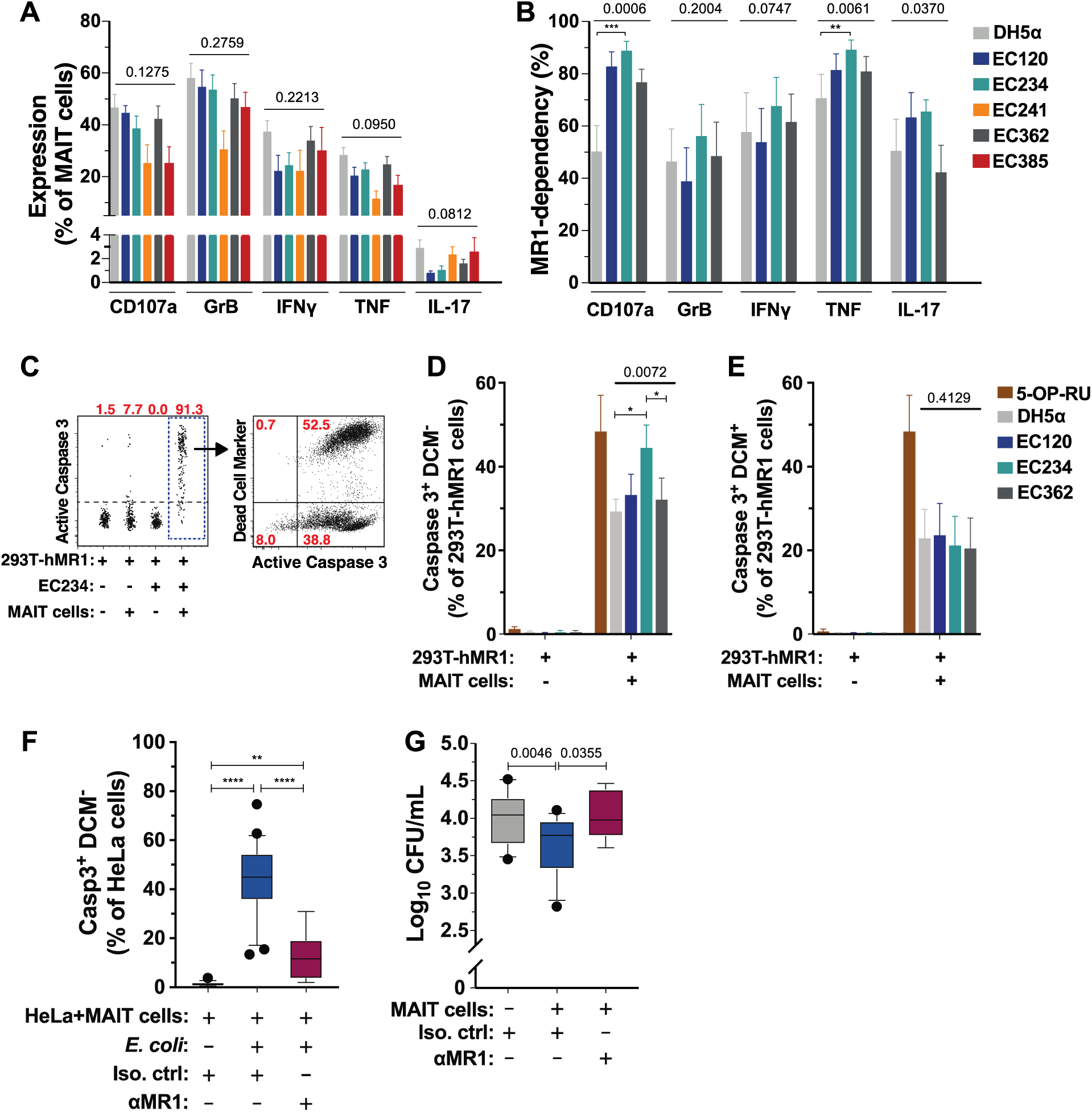
MAIT cells respond to and control CREC. (A) Expression of CD107a, GrzB, IFNγ, TNF, and IL-17A in MAIT cells stimulated for 24 h with E. coli strains DH5α, EC120S, and the carbapenem-resistant strains EC234, EC241, EC362 and EC385 (n=5 (EC241, EC385), 7 (EC120S), 16 (DH5α, EC234, EC362)). (B) MR1-dependency of effector protein and cytokine production by MAIT cells stimulated with indicated strains. MR1-dependency was calculated as previously described (49) (n=7) (C) Representative flow cytometry plots of caspase 3 activation and apoptosis in 293T-hMR1 cells alone, 293T-hMR1 cells infected with EC234, or co-culture with MAIT cells with or without EC234 for 24 h. (D, E) Caspase 3 activation and apoptosis in 293T-hMR1 cells alone or co-cultured with MAIT cells in the presence of 5-OP-RU, DH5α, EC120S, EC234, or EC362 (n=9). (F,G) Caspase 3 activation (F) and bacterial loads (G) in HeLa cells infected with EC241 co-cultured with MAIT cells in the presence of anti-MR1 antibody or isotype control (n=8 for EC241-infected HeLa cells+anti-MR1 mAb, n=16-25 others). Statistical significance was determined using the Kruskal-Wallis ANOVA (A) or the Friedman test (B-E) followed by Dunn’s multiple comparison test, and mixed-effects analysis followed by Dunnett’s multiple comparison test (F,G). * p<0.05.

To assess the capacity of MAIT cells to kill CREC-pulsed cells, the 293T cell line over-expressing human MR1 (293T-hMR1) was used as target cells as previously described [24]. MAIT cells induced caspase 3 activation and cell death in 293T-hMR1 cells pulsed with the CREC EC234 or EC362 strains, or with the drug-sensitive EC120S (Fig. 3D and E), although at lower magnitudes compared to 293T-hMR1 cells pulsed with the synthetic antigen 5-OP-RU.

To investigate whether MAIT cells can mediate antimicrobial activity against CREC strains, we selected strain EC241 because it is the CREC strain with the most efficient entry into HeLa cells (fig. S3M). MAIT cells induced caspase 3 activation in HeLa cells infected with strain EC241 in an MR1-dependent manner (Fig. 3F), consistent with results using the drug-sensitive strain EC120S (Fig. 1). Furthermore, MAIT cells significantly reduced the CREC bacterial loads within infected cells in an MR1-dependent fashion (Fig. 3G). Taken together, MAIT cells recognize CREC, mediate killing of infected cells, and reduce bacterial loads in an MR1-dependent manner.

### MAIT cells secrete high levels of cytolytic proteins and mediate antimicrobial activity against CREC in the surrounding milieu

We next investigated whether MAIT cells release cytolytic proteins into the surrounding milieu in response to MR1-restricted cognate recognition of cells infected with a riboflavin synthesis-competent strain of *E. coli*. MAIT cells secreted high levels of GrzA, GrzB, and Gnly into the supernatants already after a short 3 h stimulation with *E. coli*-infected cells (Fig. 4A). Because some cytolytic proteins have direct antimicrobial activity [28, 29], the MAIT cell secretome ability to inhibit free-living *E. coli* growth was investigated. To collect bacteria-free MAIT cell secretome following TCR-stimulation, 293T-hMR1 cells were used as antigen-presenting cells and pulsed with the synthetic bacterial riboflavin antigen 5-OP-RU and then co-cultured with MAIT cells (fig. S4A). Live *E. coli* strains were then incubated with supernatants obtained from the co-culture of MAIT cells with 5-OP-RU-pulsed 293T-hMR1 cells (fig. S4A). MAIT cells strongly degranulated their cytolytic protein content following MR1-restricted TCR triggering, and these proteins accumulated at high levels in the supernatants after 24 h of co-culture (fig. S4B and C). To assess whether MAIT cell secretomes induce bacterial damage, the cell-impermeable nucleic acid dye SYTOX Green was used [30] (fig. S4D). Bacteria were pre-stained with the cell-permeable nucleic acid dye SYTO 62 to allow their detection by flow cytometry (Fig. 4B). In the presence of MAIT cell supernatants, the CREC strains EC234 and EC362 suffered an increase in membrane permeability, as seen by the significant increase of SYTOX Green and SYTO 62 staining when compared to the control supernatants (Fig. 4B and C, fig. S4E and F). Finally, the bacterial counts of the *E. coli* strains exposed to the MAIT cell secretomes were significantly reduced, although not totally suppressed (Fig. 4D). Taken together, these results indicate that the MAIT cell secretomes display antimicrobial activity against *E. coli* by markedly increasing bacterial cell permeability.

**Fig. 4.**
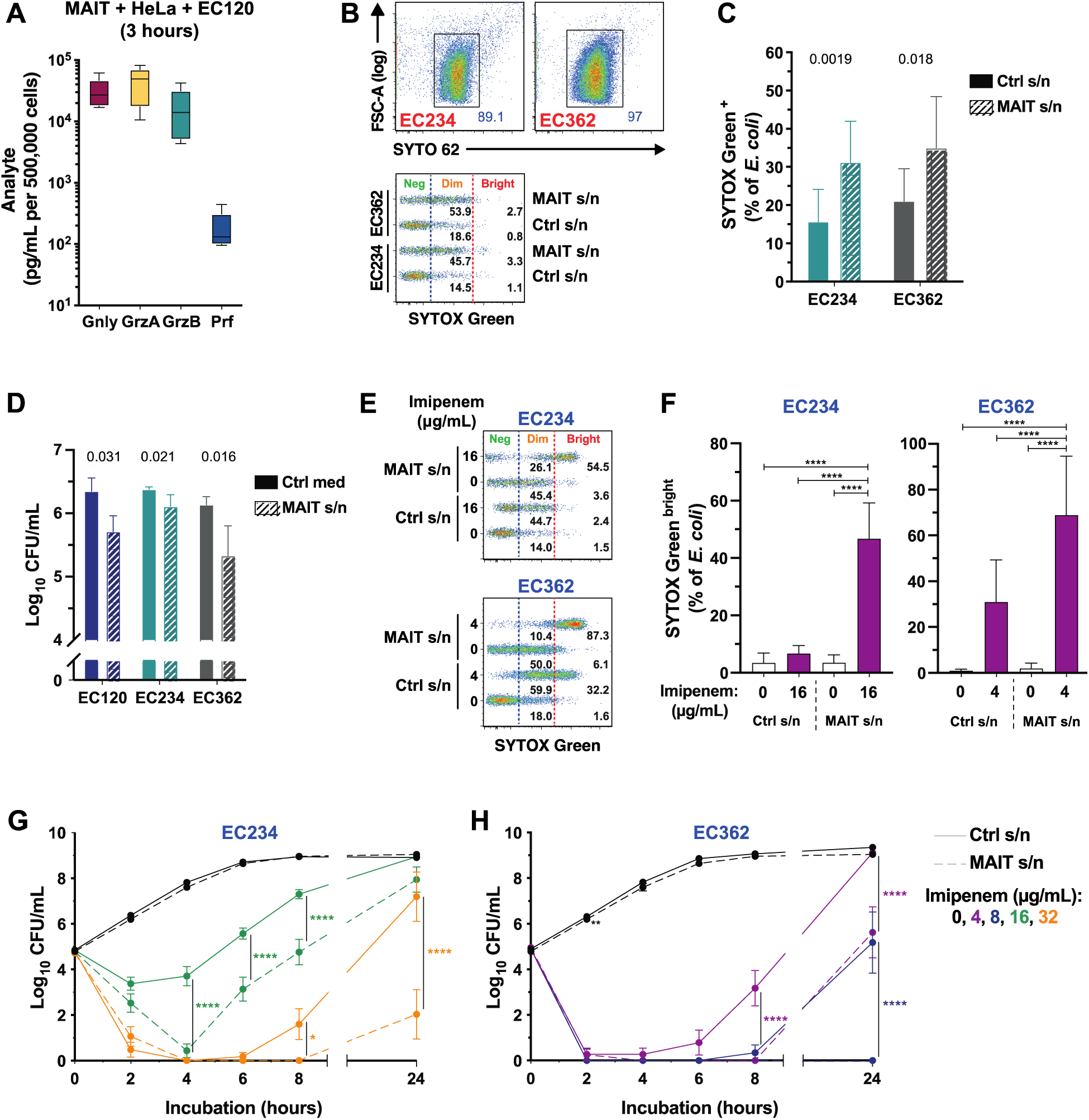
MAIT cell secretomes mediate antimicrobial activity against CR-E-coli. (A) Concentration of cytolytic proteins in the supernatant of MAIT cells following 3 h co-culture with E. coli EC120S-infected HeLa cells (n=6). (B) Gating strategy for E. coli identification and quantification of SYTOX Green by flow cytometry. (C) SYTOX green levels (n=9) and (D) bacterial counts of various E. coli strains after incubation with MAIT cell or control supernatants for 2 h (n=6 (EC120S), others n=12). (E, F) Levels of SYTOX Green^bright^ in E. coli EC234 and EC362 in the presence of MAIT cell or control supernatants supplemented with imipenem for 2 h (n=10) or (G, H) the live bacterial counts over 24 h (n=15-17 (EC234), 11-13 (EC362)). Statistical significance was determined using paired t-test (C), Wilcoxon’s signed-rank test (D), RM one-way ANOVA with Dunnett’s post-hoc test (F), and mixed-effects analysis with Sidak’s post-hoc test (G, H). Significant differences in bacterial counts cultured in control vs. MAIT cell supernatants at indicated time-points and imipenem concentrations in (G) and (H) were indicated by asterisks (**** p<0.0001, ** p<0.01, * p<0.05). S/n, supernatant.

### MAIT cell secretomes enhance the bactericidal activity of carbapenems against CREC

Following the observation that MAIT cell secretomes can directly alter bacterial membrane permeability (Fig. 4B-D), we asked whether the MAIT cell secretome could enhance the bactericidal activity of carbapenems against CREC strains. To test this, CREC strains EC234 and EC362 were incubated in the presence of MAIT cell supernatants with titrated concentrations of the carbapenem antibiotics imipenem, ertapenem, and meropenem. Greatly enhanced permeability and damage to the bacteria was evident by the appearance of bacteria stained with high intensities for SYTOX Green (SYTOX Green^bright^), in comparison with bacteria cultured in either MAIT cell supernatants or carbapenem antibiotics alone (Fig. 4E and F, fig. S4G,H). Fluorescence intensity of SYTO 62 was also increased in bacteria treated with a combination of MAIT cell secretome and imipenem (fig. S4I and J).

Next, effects on the live bacterial counts were evaluated using a time-kill method [31]. Notably, MAIT cell secretomes in combination with imipenem killed the highly carbapenem resistant *E. coli* strain EC234 (minimum inhibitory concentrations (MICs) to all carbapenems ≥32μg/mL; Table S2), as well as the XDR strain EC362 harboring further resistance to colistin (Table S1). In both cases, imipenem concentrations were well below the strains’ respective MICs (Fig. 4G and H). Growth rates of strains EC234 and EC362 cultured in MAIT cell supernatants including titrated concentrations of imipenem were significantly slower compared to their respective controls (fig. S5A and B). These growth delays occurred at imipenem concentrations up to 4-fold lower than the MIC for EC234, and up to 16-fold lower than the MIC for EC362 (fig. S5A and B). Moreover, the MAIT cell secretome decreased the imipenem concentrations required to fully suppress the growth of strains EC234 and EC362 by at least 2- and 4-fold, respectively (fig. S5C). In the presence of imipenem, the antimicrobial activity of MAIT cell secretomes appeared to be superior to that of non-MAIT Vα7.2^-^ T cells from the same donors (fig. S5D). Finally, we evaluated whether the antimicrobial activity was derived from the MAIT cells themselves, or from the dying 293T-hMR1 cells used as antigen-presenting cells. Firstly, we showed that supernatants of Vα7.2 bead-purified and expanded MAIT cells cultured without 293T-hMR1 cells still displayed antimicrobial activity in the presence of imipenem against the XDR strain EC362 (fig. S5E and F). Secondly, blocking the apoptosis of 293T-hMR1 cells caused by MAIT cell cytotoxicity with the pan-caspase inhibitor Pro-VAD-FMK, caused no significant loss of antimicrobial activity in the presence of imipenem against strains EC234 and EC362 (fig. S5G and H). Taken together, these findings suggest that there is a synergy between MAIT cell secretome antimicrobial activity and carbapenems, thereby enhancing the *in vitro* bactericidal activity of carbapenems against CREC strains.

### MAIT cell secretome antimicrobial activity correlates with the levels of secreted cytolytic proteins

We next re-visited the time-kill data (Fig. 4G and H), and analyzed possible associations between the amounts of cytolytic proteins present in the MAIT cell supernatants and the antimicrobial activity when combined with imipenem. In the bacterial cultures where growth was detected, there was significantly less Gnly and GrzB in the MAIT cell secretome (Fig. 5A and B). However, no significant difference in GrzA and Prf levels was observed (Fig. 5A and B). Moreover, there were negative correlations between viable bacterial counts and Gnly or GrzB levels within the MAIT cell secretomes (Fig. 5C-F). However, no correlations with the levels of GrzA or Prf were observed (fig. S6A-D). Interestingly, Gnly levels in the supernatants correlated positively with the length of the lag-phase for strains EC234 and EC362 in the presence of imipenem (fig. S6E and F). Furthermore, the MAIT cell secretomes that contained the highest concentrations of Gnly had significantly greater and longer inhibition on EC234 and EC362 (fig. S6G). Taken together, these findings suggest that cytolytic proteins, particularly Gnly and GrzB, may significantly contribute to the MAIT cell secretome antimicrobial activity against CREC in the presence of imipenem.

**Fig. 5.**
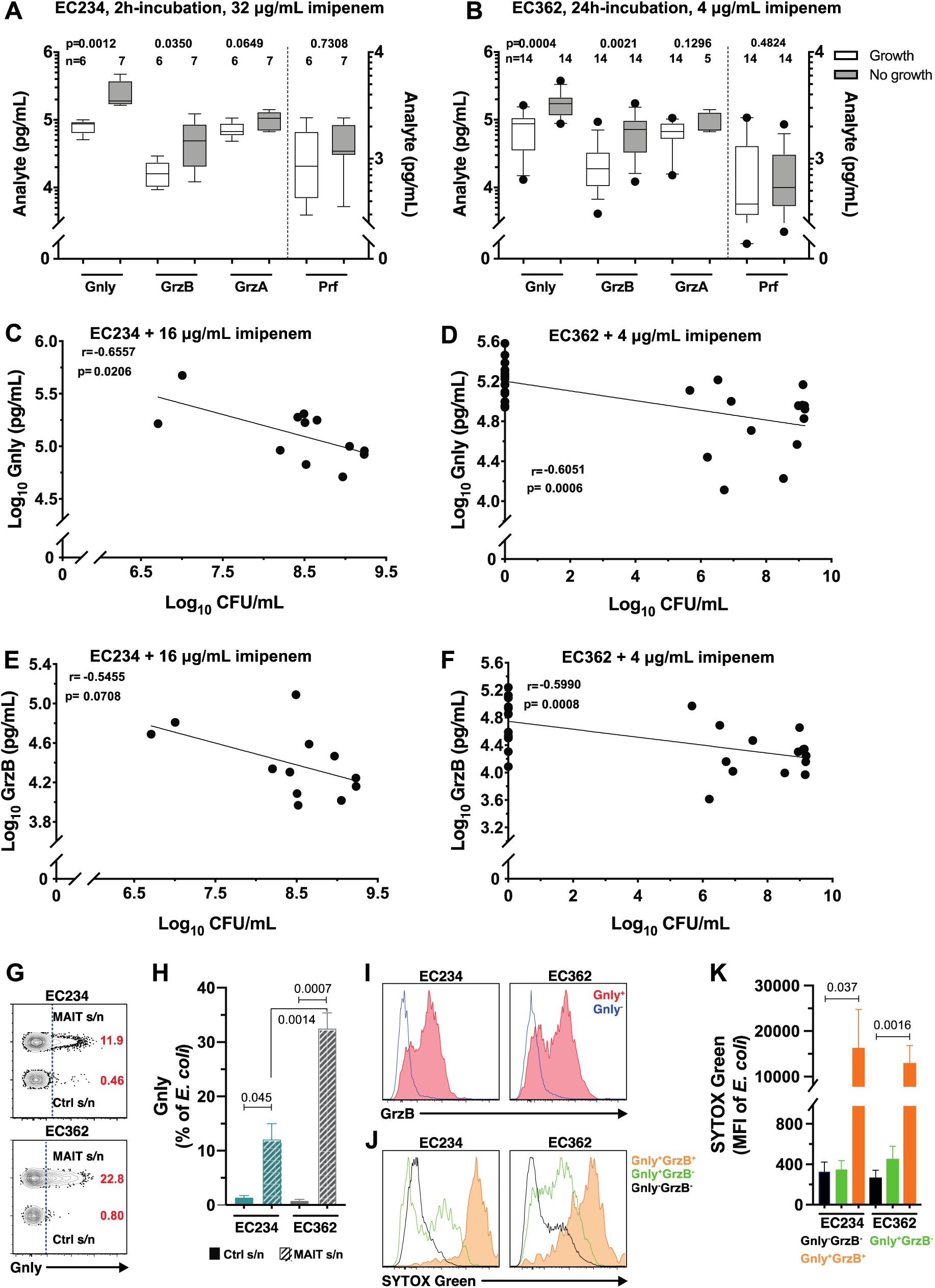
Cytolytic proteins contribute to the antimicrobial activity of MAIT cell secretomes. (A,B) Concentration of cytolytic proteins in the MAIT cell supernatants spiked with imipenem in the E. coli EC234 and EC362 cultures where growth was detected or not at the indicated time-points. (C, D) Correlation between Gnly and (E, F) GrzB levels in the MAIT cell supernatants and E. coli EC234 and EC362 bacterial loads after 24 h incubation in the MAIT cell supernatants spiked with imipenem (n=12 (EC234), 28 (EC362)). (G) Gating strategy of Gnly flow cytometry staining and (H) proportion of Gnly in E. coli EC234 and EC362 following 30 min incubation with MAIT cell or control supernatants (n=4-6). (I) Representative histograms of GrzA, GrzB, Prf, and (J) SYTOX Green staining and (K) intensity in E. coli EC234 and EC362 following 30 min incubation in MAIT cell or control supernatants (n=4-6). Statistical significance was determined using Mann-Whitney’s test (A, B), paired t-test for intra-strain or unpaired t-test for inter-strain analyses (H), and Friedman’s test with Dunn’s post-hoc test (K). Correlations were calculated using the Spearman test.

### Gnly and GrzB contribute to the antimicrobial activity of MAIT cell secretomes and potentiate the bactericidal action of carbapenems against CREC

Finally, the functional contribution of different cytolytic proteins to the MAIT cell secretome antimicrobial activity was examined. To enable this, the assay dynamic ranges was enhanced by using strain EC362, which was more sensitive to the inhibitory effects of the MAIT cell secretomes (Fig. 4H), and a lower amount of imipenem (2 μg/mL), to reduce its residual inhibitory effects. Blocking MAIT cell degranulation using EGTA + Mg^2+^ (fig. S6H) nearly completely abolished the MAIT cell secretome-mediated bacterial inhibition (fig. S6I), whereas blocking GrzB activity alone had no observable effect (fig. S6I). Similarly, blocking the activity of IFNγ, TNF, or IL-17A during the MAIT cells-293T-hMR1 cells co-culture period had no significant effect on the antimicrobial activity of harvested supernatants (fig. S6I). Importantly, specific depletion of Gnly from the supernatants (fig. S6J) significantly reduced the MAIT cell secretome inhibition of bacterial growth (fig. S6K), as well as the capacity to induce bacterial cell permeability (fig. S6L). Gnly was detected intracellularly by flow cytometry in both EC234 and EC362 bacterial cells incubated with MAIT cell supernatants (Fig. 5G and H, fig. S7A). GrzB was also detected, but mostly in Gnly-containing bacteria (Fig. 5I), while no GrzA or Prf was detected in bacterial cells (Fig. 5I, fig. S7A and B). Notably, bacterial cells containing both Gnly and GrzB had higher membrane permeability than those containing only Gnly (Fig. 5J and K). Short-term concomitant treatment with imipenem (30 min) did not increase Gnly levels inside bacterial cells despite the increase in bacterial permeability and damage (fig. S7C). This suggests that the increase in bacterial death following incubation with MAIT cell supernatants and imipenem (Fig. 4H), was possibly caused by the higher imipenem penetration into bacterial cells following MAIT cell-mediated increase in bacterial cell permeability. Collectively, these findings indicate that Gnly and GrzB contribute to the antimicrobial activity of MAIT cell secretomes and potentiate the bactericidal action of carbapenem antibiotics against CREC (Fig. 6).

**Fig. 6.**
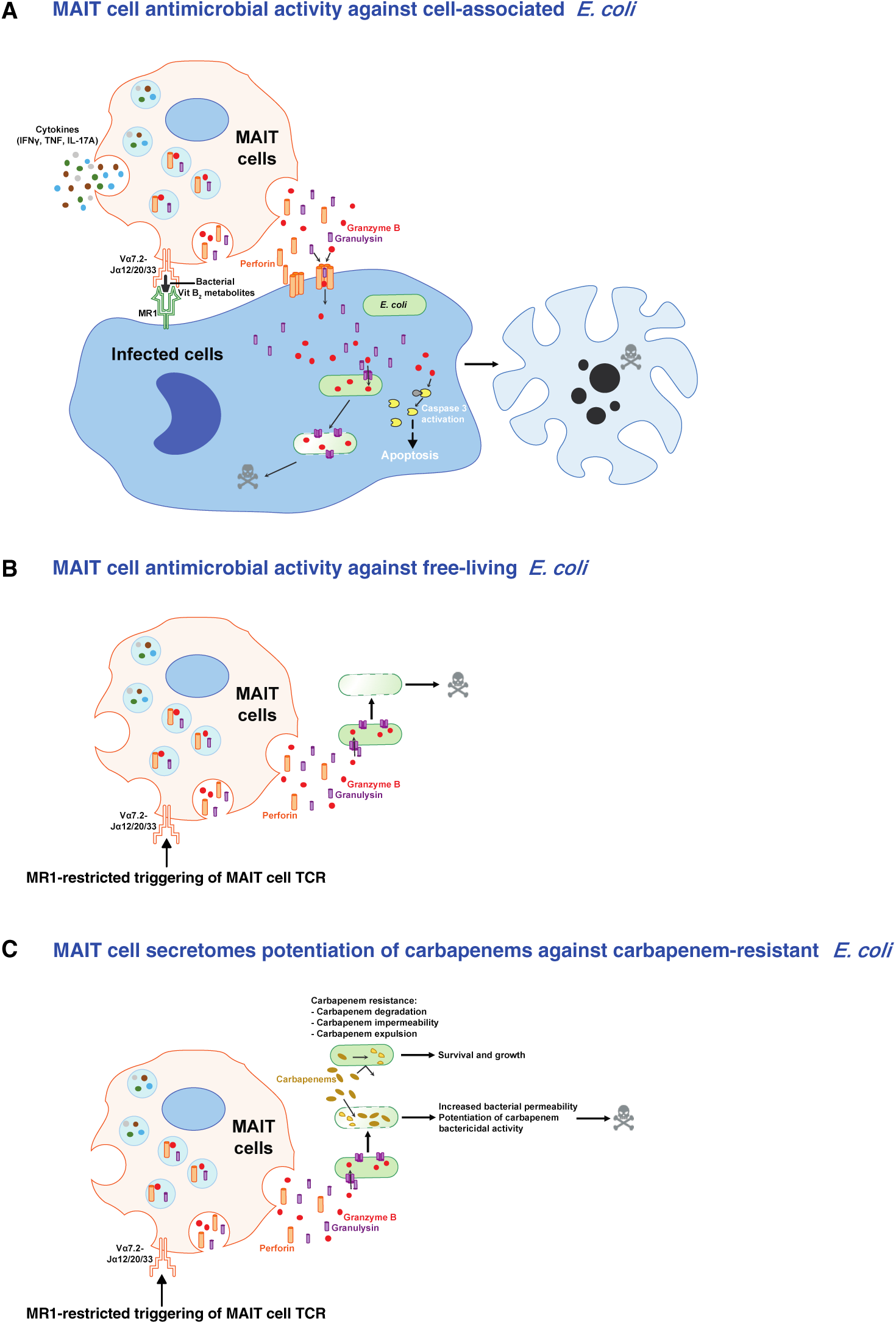
A model of MAIT cell antimicrobial activity against cell-associated (A) and free-living (B) drug-sensitive and carbapenem-resistant E. coli, and (C) MAIT cell secretome potentiation of carbapenem killing activity against carbapenem-resistant E. coli strains.

## Discussion

Despite a considerable body of evidence that MAIT cells play a protective role in various bacterial infections, it is not known whether MAIT cells directly inhibit bacterial growth nor what mechanism is operating in mediating antimicrobial activity. It is also unknown whether MAIT cells have antimicrobial activity against drug-resistant bacteria. Here, we show that human MAIT cells have direct antimicrobial activity against both cell-associated and free-living *E. coli* primary clinical isolates, including CREC strains that are extensively resistant to carbapenems. This antimicrobial activity depends on the TCR-mediated activation of cytolytic protein secretion by MAIT cells in response to cognate recognition of cells that were infected or have taken up bacteria and presented antigen. High levels of cytolytic effector proteins in the MAIT cell secretome directly damage free-living CREC bacterial cells by increasing their permeability. Strikingly, MAIT cell secretomes restore the bactericidal activity of carbapenems against free-living CREC clinical isolates *in vitro*. The levels of cytolytic proteins secreted by MAIT cells, in particular Gnly and GrzB, correlate with the degree of bacterial killing and the potentiating effect of carbapenem bactericidal activity. Finally, blocking experiments identified Gnly as an important component of the antimicrobial activity of MAIT cell secretomes. Altogether, these results demonstrate potent antimicrobial activity of MAIT cells against both cell-associated and free-living *E. coli*, and show that MAIT cells can act to restore the antibiotic effect of carbapenems against CREC.

MAIT cells promote antimicrobial effects through multiple mechanisms, including killing of infected cells, induction of nitric oxide production, and orchestrating downstream effector cell responses [14, 25, 32–37]. Here, our findings indicate that both Gnly and GrzB expression are required for efficient MAIT cell bacterial control, independent of infected cell death and production of pro-inflammatory cytokines. Consistent with this model, a recent study revealed the existence of an adaptive CD8^+^ T cell subset co-expressing Prf, GrzB and Gnly, which can kill intracellular bacteria [38]. In the context of cytolytic protein secretion by cytotoxic CD8^+^ T cells, Gnly-mediated pore formation in bacterial cell walls allows GrzB to penetrate into bacterial cytoplasm and cleave oxidative stress defense enzymes and other proteins vital for bacterial metabolism. This ultimately leads to the generation of reactive oxygen species (ROS) and bacterial death [28, 29]. Interestingly, in a mouse model of *Legionella* infection, cytolytic proteins did not appear to contribute to the MAIT cell protective role [36]. However, because cytolytic proteins are structurally and functionally divergent in mammals [39] and Gnly is absent in rodents [40], prudence should be exercised in interpreting the mechanism underlying MAIT cell mediated antimicrobial activity in murine models. Collectively, our findings indicate that MAIT cells mediate antimicrobial activity through the cytolytic protein-dependent pathway, which may contribute to the protection of the human host against bacterial infections.

An important aspect of the present study concerns the differential and temporal regulation of MAIT cell cytotoxicity and antimicrobial activity. Resting MAIT cells expressed GrzA and Prf but were negative for GrzB and Gnly, with no appreciable cytotoxic or antimicrobial capacity. Following antigenic stimulation there was a rapid and strong upregulation of GrzB, accompanied by strong cytotoxicity but inefficient antimicrobial activity, and finally a gradual expression of Gnly with strong cytotoxicity and antimicrobial activity. How long these cytolytic effector proteins remain expressed by MAIT cells following antigenic stimulation and resolution of bacterial infection, and whether such temporal regulation influences MAIT cells’ protective roles of the host is unclear. We previously observed variable expression levels of Gnly in resting peripheral blood MAIT cells [2, 25]. As Gnly is similarly expressed late by activated cytotoxic T cells following activation [41], Gnly^+^ MAIT cells in blood may represent an antigen-imprinted and -experienced MAIT cell subset that may respond to bacterial infection faster. While Gnly is broadly antimicrobial, it also has strong pro-inflammatory and chemotactic activity, and has been implicated in several inflammatory diseases [40]. Thus, the strict differential and temporal regulation of cytotoxicity and antimicrobial activity may also afford some protection against MAIT cell-mediated immunopathology that can occur in certain bacterial infections [42, 43]. Interestingly, cytolytic proteins are differentially expressed during the development of human MAIT cells, with the mature subsets expressing more of these effector proteins [44]. In adult human female genital tract, oral mucosa, and placental tissues, MAIT cells express low level of GrzB at steady state [12, 45–47], suggesting that the regulation of cytolytic effector proteins in tissues may differ from that of blood MAIT cells.

CRE organisms, including CREC, have a high risk of transmission and rapid spread between patients in the hospital settings [20]. Carbapenems disrupt bacterial cell wall synthesis and the resistance to carbapenems in *Enterobacteriaceae* often involves carbapenem impermeability, efflux pumps, and expression of carbapenemases. Combined with the increasing occurrence of pan-drug resistance, the spread of CRE organisms are a rising threat to public health [20]. Few studies have considered the role of the host’s immune response in facilitating clearance of resistant populations arising during the course of antimicrobial therapy. Yet, strong host’s immune response during the course of antimicrobial chemoterapy has been proposed to limit the duration of treatment and appearance of resistant populations [48–50]. This is one possible explanation for why treatments with single antimicrobial agents most of the time are successful in healthy, immunocompetent individuals [51]. In the present study, we show that MAIT cells recognize and respond to CREC-infected cells, and mediate efficient killing of infected cells and rapid control of bacterial loads. Our findings that MAIT cells retain their ability to mediate antimicrobial activity against resistant bacteria, implies that MAIT cells play a role in defense not only against drug-sensitive bacteria, but also help fend off drug-resistant bacteria in hosts that have functional MAIT cells. Consistent with this hypothesis, individuals with co-morbidities that are frequently linked with low MAIT cell numbers and poor antimicrobial functionality [52–55], are more susceptible to infections caused by drug-resistant bacterial pathogens, incl. CRE [56–59]. This suggests the potential importance of MAIT cell antimicrobial effector response in the host’s defense against drug-resistant bacterial pathogens. It is thus tempting to speculate that it may be possible to prevent CRE infections among at-risk individuals by restoring MAIT cell numbers and functionality through vaccinations using MR1 ligands [37].

Finally, and strikingly, high levels of cytolytic effector proteins secreted by MAIT cells in response to MR1-restricted cognate recognition of bacterial riboflavin antigens potentiate the bactericidal activity of carbapenems against CREC primary clinical isolates. This restoration of carbapenem activity *in vitro* is likely due to increased bacterial membrane permeability mediated by high levels of Gnly, allowing entry of lethal amounts of carbapenems into the bacteria. Consistent with this hypothesis, we detected the presence of Gnly in bacterial cells exposed to MAIT cell secretomes, and this was associated with increased bacterial permeability and damage. Gnly is a cationic, saposin-like antimicrobial protein that binds to and forms pores on cholesterol-poor bacterial membranes, and kill bacteria by disrupting membrane permeability, including those of *E. coli* [30, 60–62]. Previous studies showed that GrzB enters the bacterial cytoplasm *via* pores formed by Gnly and induces oxidative stress damage [28, 29]. Consistent with this, we detected intracellular GrzB in bacterial cells incubated with MAIT cell secretomes, but only when Gnly was also present. The dual presence of GrzB and Gnly further increased bacterial permeability and damage. Thus, GrzB contributed to the antimicrobial effector function of MAIT cell secretomes and their potentiation of carbapenem killing activity. It is tempting to hypothesize that MAIT cell secretome antimicrobial activity is not only able to control bacterial growth, but also able to disarm carbapenem impermeability mechanisms of resistance in CREC by increasing carbapenem access to bacterial intracellular space. In conclusion, the findings in the present study indicate that MAIT cell cytolytic properties are an important component of their antimicrobial effector function and enable enhancement of carbapenem killing activity against carbapenem-resistant *E. coli* strains. The ability of MAIT cell antimicrobial effector function to overcome drug resistance suggest that they may participate in clearance of bacteria acquiring *de novo* resistance during the course of antibiotic therapy.

### Blood processing and MAIT cell expansion

Peripheral blood was collected from healthy donors recruited at the apheresis unit, Bloodbank@HSA, Health Services Authority, Singapore. Written informed consent was obtained from all donors and ethical approval was obtained from the National University of Singapore Institutional Review Board (NUS-IRB reference codes B-15-088 and H-18-029). Peripheral blood mononuclear cells (PBMCs) were isolated by standard Ficoll-Histopaque density gradient separation (Ficoll-Histopaque Premium; GE Healthcare). After isolation, PBMCs were cryopreserved in liquid nitrogen until further use, or immediately used for MAIT cell purification or activation assays.

MAIT cells were expanded using two different protocols in this study. For the first set of expansion, MAIT cell were isolated from freshly-isolated PBMCs by positive selection using the 5-(2-oxopropylideneamino)-6-D-ribitylaminouracil (5-OP-RU)-loaded human MR1 tetramer-PE using magnetic-activated cell sorting (MACS) and anti-PE microbeads (Miltenyi). MAIT cells (>95% purity) were cultured 6-7 days in serum-free and xeno-free ImmunoCult-XF T cell expansion medium (STEMCELL Technologies) in the presence of 10 ng/mL recombinant human (rh)IL-7 (R&D Systems), 1:100 ImmunoCult human CD3/CD28/CD2 T cell activator (STEMCELL Technologies), 50 μg/mL gentamicin (Gibco), and 100 μg/mL normocin (Invivogen). For the second set of expansion, cryopreserved PBMCs were thawed and cultured in ImmunoCult-XF T cell expansion medium supplemented with 8% (v/v) xeno-free CTS Immune Cell Serum Replacement (ThermoFisher Scientific), 5 ng/mL rhIL-2 (Peprotech), 10 ng/mL rhIL-7, 50 μg/mL gentamicin, and 100 μg/mL normocin. PBMCs were stimulated with 10 nM 5-OP-RU on day 0, 5, and 10, and the culture media was replenished every 2-3 days. On day 11, viable cells were isolated by Ficoll-Histopaque density gradient centrifugation. On day 15-17 cells were checked for MAIT cell purity and numbers by flow cytometry. Cells were immediately used for subsequent assays when MAIT cell purity > 70 %.

### Bacterial cultures

The *E. coli* strains 1100-2 and BSV18 were obtained from the Coli Genetic Stock Center, Yale University; the DH5α strain was obtained from New England Biolab. Carbapenem-resistant isolates were identified from the Singapore General Hospital microbiological database, and retrieved from the archived bacteria repository. Further details on strain identification and determination of resistance and susceptibility profiles can be found in the Supplementary Materials section.

For MAIT cell stimulation assays, all *E. coli* strains were grown overnight at 37°C in Luria (lysogeny) broth (LB) with shaking as described [63]. Overnight cultures of *E. coli* were then subcultured 10-fold in LB and incubated at 37 °C with shaking until OD_600_ = 0.5. In selected experiments, growth curves were monitored by reading absorbance at 600 nm in a microplate reader with discontinuous shaking for 18 h at 37 °C (Cytation 5, BioTek Instruments).

### MAIT cell functional and antimicrobial activity assays

MAIT cells within bulk PBMCs, identified as MR1-5-OP-RU^+^ Vα7.2^+^ CD161^hi^ CD3^+^ T cells, were activated using formaldehyde-fixed *E. coli* strains as indicated for 24 h as previously described [24]. In selected experiments, 20 μg/mL MR1 blocking mAb (26.5, Biolegend) or IgG2a isotype control (MOPC-173, Biolegend) were used (*22*). In all activation assays, monensin (Golgi Stop, BD Biosciences) were added for the last 6 h of incubation.

MAIT cell cytotoxicity assay was performed as previously described [24, 25]. Briefly, human 293T cells stably transfected with human MR1 (293T-hMR1) were incubated in complete RPMI medium with formaldehyde-fixed *E. coli* at the microbial dose of 30 for 3 h before the addition of expanded MAIT cells at the effector to target cell ratio 5:1. 2 nM 5-OP-RU was added as a positive control for MAIT cell killing of target cells. Anti-CD107a-BUV395 at 1:200 was added at the beginning of the assay to detect MAIT cell degranulation. After 24 h of co-culture, cells were stained to detect target cell apoptosis using anti-active caspase 3 mAb (BD Biosciences). MAIT cell antimicrobial activity assay was performed as described [64]. Briefly, the bacteria sub-cultures were resuspended in RPMI without serum and antibiotics (ASF-RPMI). Adherent target epithelial cell lines were infected with live *E. coli* for 3 h at a microbial dose of 30 for strain EC120S and 3 for strain EC241 at 37 °C / 5% CO_2_. Infected cells were washed extensively with complete RPMI medium supplemented with 200 μg/mL gentamicin (Gibco) and further incubated for 1 h at 37 °C / 5 % CO_2_ to kill extracellular bacteria, then washed extensively with ASF-RPMI. Expanded MAIT cells were labelled with 1 μM CellTrace Violet (CTV) dye (Thermo Fisher Scientific) before co-cultured with HeLa cells at an E:T ratio of 5:1 and incubated for 3 h at 37 °C / 5 % CO_2_. For the live bacteria enumeration, a duplicate set of experimental wells were done in parallel. Supernatants from the first set were collected, followed by adherent cell lysis with 0.1% (v/v) Triton-X for 10 min at room temperature. Equivalent volume of LB broth were added to lysates and plated onto LB agar plates in duplicates, incubated at 37 °C for 18 to 24 h, and counted visually. For the second set, anti-CD107a-BUV395 (1:200) was added into the culture medium at the start of the assay to assess MAIT cell degranulation. Cell-free supernatants were harvested, snap frozen in liquid nitrogen, and stored at -80 °C until further use. Adherent cells were harvested using trypsin-EDTA (Gibco) and stained for flow cytometry as indicated.

In selected experiments, MAIT cells or infected HeLa cells were treated with various pharmacological inhibitors or mAbs before use in assays. Briefly, MAIT cells were pre-incubated for 1 h with 5 mM EGTA (Bioworld) supplemented with 1 mM MgCl_2_ (Sigma-Aldrich), then diluted to 1 mM EGTA with 1 mM MgCl_2_ in-assay, or with 10 μM nafamostat mesylate (Sigma-Aldrich), or 100 μM of the GrzB inhibitor II Ac-IETD-CHO (Merck) before co-culture with HeLa cells. HeLa cells were pre-treated for 1 h before addition of MAIT cells with 10 μM CAS-BIND Pro Pan Caspase Inhibitor (Pro-VAD-FMK) (Vergent Bioscience). For antibody-blocking experiments, 20 μg/mL anti-MR1 or IgG2a isotype control was added on HeLa cells 1 h prior to co-culture, or 10 μg/mL mAb to IFNγ (B27; Biolegend), TNF (MAb1; Biolegend), IL-17A (eBio64CAP17; Invitrogen) or IgG1 isotype control (MOPC-21; Biolegend) 15 min before co-culture.

### Granzyme B activity detection

Granzyme B activity was measured with GranToxiLux PLUS! (GTL) Kit (Oncoimmunin, Inc). Briefly, Hela cells were trypsinized and infected with *E. coli* as per MAIT cell antimicrobial assay and MAIT cells were co-cultured with HeLa cells at E:T ratio of 5:1. Co-cultured cells were centrifuged immediately and resuspended in GrzB Substrate solution. The co-culture was done for 1 h at 37 °C and cells were washed in the GTL Wash Buffer. Cells were stained with viability dye and immediately analyzed by flow cytometry.

### Preparation of MAIT cell supernatants

293T-hMR1 cells were plated in a 96-well flat bottom plate at 37°C / 5 % CO_2_ in complete RPMI medium. After overnight incubation, culture medium was replaced with the ImmunoCult medium and 2 nM 5-OP-RU was added into each well for 2 h. Expanded MAIT cells were then further purified using the Vα7.2 beads as described (*22*) (MAIT cell purity >98%) and co-cultured with the 5-OP-RU-pulsed 293T-hMR1 cells at an E:T ratio of 10:1. Supernatants from the co-culture wells or 5-OP-RU-pulsed 293T-hMR1 cell control wells were collected 24 h after co-culture, clarified, and snap-frozen in liquid nitrogen until further use. The cytokine and cytolytic protein contents of the supernatants were measured using the LEGENDplex human CD8/NK cell panel (Biolegend) as described [2].

### MAIT cell secretome antimicrobial activity assay

Overnight bacteria subcultures were washed with PBS and resuspended in MAIT cell or control supernatants at 10^5^ CFU/mL and incubated at 37 °C in flat-bottom 96-well plates without shaking for 24 h. To enumerate live bacteria, bacterial suspensions were harvested at various time points as indicated, serially diluted in LB broth, plated in triplicates on LB agar plates, and incubated at 37 °C for 18 to 24 h and counted visually. In selected experiments, MAIT cell and control supernatants were spiked with imipenem monohydrate (Sigma-Aldrich) at various concentrations as indicated. A three log_10_ reduction in bacterial counts from the baseline population over 24 h was considered as bactericidal.

To assess bacterial damage, the cell-impermeable SYTOX Green nucleic acid stain (Thermo Fisher Scientific) was added at 20 μM during the 2 h-incubation of 10^6^ CFU/mL *E. coli* with MAIT cell or control supernatants, with or without carbapenems at the indicated concentrations. Dead bacteria controls were prepared by incubating the bacteria in 70% (v/v) ethanol for 30 min at room temperature, washed, then incubated with control medium supplemented with Sytox Green. To allow identification of the bacteria versus debris by flow cytometry, the bacteria were stained with SYTO 62 red fluorescent nucleic acid stain (Thermo Fisher Scientific; 2.5 μM) during the last 15 min at 37 °C. Bacteria were then fixed with 1% formaldehyde for 20 min at 4 °C just prior to FACS acquisition. In selected experiments, to detect the intracellular cytolytic proteins in bacterial cells, 10^5^ CFU/mL of *E. coli* were cultured with control or MAIT cell supernatants in the presence of 20 μM SYTOX Green for 30 min at 37 °C without shaking. During the last 15-min of incubation, fluorochrome-conjugated mAbs against Gnly, GrzA, GrzB, and Prf (Table S3), as well as SYTO 62 nucleic acid dye were added to the culture. Bacteria were then washed, fixed, and permeabilised with BD Cytofix/Cytoperm buffers (BD Biosciences), then re-stained with the same mAb cocktail for 30 min at 4 °C and washed once prior to FACS acquisition.

### Flow cytometry analysis

Cell surface and intracellular staining for cytokines, cytotoxic molecules, and active caspase 3 were performed as previously described [2]. Staining with the MR1 5-OP-RU and MR1 6-FP tetramers was performed for 40 min at room temperature (RT) [7] before proceeding to the surface and intracellular staining with other mAbs (Table S3.) Samples were acquired on an LSRFortessa flow cytometer (BD Biosciences) equipped with 355, 405, 488, 561, and 639 nm lasers. Single-stained polystyrene beads (BD Biosciences) and the compensation platform in FACSDiva v. 8.0.1 (BD Biosciences) or FlowJo software v. 9.9 and 10.5 (TreeStar) were used for compensation.

### Statistical analysis

Statistical analyses were performed using Prism software v.8.3.0 (GraphPad). Data sets were first assessed for data normality distribution. Data presented as heatmap shows the mean, whereas data presented as line or bar graphs with error bars represent the mean and standard error. Box and whisker plots show median, the 10^th^ to 90^th^ percentile, and the interquartile range. Statistically significant differences between samples were determined as appropriate using the unpaired t-test or Mann-Whitney’s test for unpaired samples, and the paired t-test or Wilcoxon’s signed-rank test for paired samples. The Kruskal-Wallis one-way analysis of variance (ANOVA), the Friedman test, ordinary ANOVA, the repeated-measures (RM) one-way ANOVA, or mixed-effects analysis followed by the appropriate post-hoc test as indicated was used to detect differences across multiple samples. Correlations were assessed using the Spearman rank correlation. Two-sided p-values < 0.05 were considered significant.

## Supplementary Materials

Supplementary Materials and Methods

Fig. S1. MAIT cells killed E. coli-infected cells and suppressed bacterial loads.

Fig. S2. Expression of cytolytic proteins in MAIT cells is temporally regulated.

Fig. S3. MAIT cells responses to stimulation with CREC clinical strains.

Fig. S4. Antimicrobial activity of MAIT cell secretomes.

Fig. S5. MAIT cell secretomes restore imipenem bactericidal activity against CREC strains.

Fig. S6. Relationship between MAIT cell-derived cytolytic proteins and antimicrobial activity of MAIT cell secretomes.

Fig. S7. Detection of cytolytic proteins inside bacterial cells. Table S1. The *E. coli* clinical isolates used in this study.

Table S2. The minimum inhibitory concentrations (MIC) of the *E. coli* clinical isolates. Table S3. Flow cytometry-based antibodies and reagents used in the study.

## Acknowledgments

We thank Dr. Ted Hansen for the kind gift of 293T-hMR1 cell line. The MR1 tetramer technology was developed jointly by Dr. James McCluskey, Dr. Jamie Rossjohn, and Dr. David Fairlie; and the material was produced by the NIH Tetramer Core Facility as permitted to be distributed by the University of Melbourne.

## Author contributions

CB, WRS, MYG, TKP performed the experiments. CB, WRS, EL analysed the data. CB, WRS, EL designed the experiments. JQMT, JYWM, DPF, IKHP, THK, LFW, ALHK provided critical reagents. JKS, EL conceived the study. ALHK, JKS, EL managed the study. CB, WRS, JKS, EL wrote the paper, which was approved by all authors.

## Data and materials availability

All data associated with this study are present in the paper and/or the Supplementary Materials. The E. coli clinical isolates described in the study are available under a material transfer agreement, and the whole gemome sequence data set has been deposited to the NCBI Bioproject (PRJNA577535).

## References and Notes

1. Godfrey DI, Koay HF, McCluskey J, Gherardin NA. The biology and functional importance of MAIT cells. Nat Immunol. 2019;20(9):1110–28. Epub 2019/08/14. doi: 10.1038/s41590-019-0444-8. PubMed PMID: 31406380.

2. Dias J, Boulouis C, Gorin JB, van den Biggelaar R, Lal KG, Gibbs A, et al. The CD4(-)CD8(-) MAIT cell subpopulation is a functionally distinct subset developmentally related to the main CD8(+) MAIT cell pool. Proc Natl Acad Sci USA. 2018;115(49):E11513–E22. Epub 2018/11/18. doi: 10.1073/pnas.1812273115. PubMed PMID: 30442667; PubMed Central PMCID: PMCPMC6298106.

3. Kurioka A, Jahun AS, Hannaway RF, Walker LJ, Fergusson JR, Sverremark-Ekström E, et al. Shared and Distinct Phenotypes and Functions of Human CD161++ Vα7.2+ T Cell Subsets. Front Immunol. 2017;8. doi: 10.3389/fimmu.2017.01031.

4. Treiner E, Duban L, Bahram S, Radosavljevic M, Wanner V, Tilloy F, et al. Selection of evolutionarily conserved mucosal-associated invariant T cells by MR1. Nature. 2003;422(6928):164–9. Epub 2003/03/14. doi: 10.1038/nature01433. PubMed PMID: 12634786.

5. Mondot S, Boudinot P, Lantz O. MAIT, MR1, microbes and riboflavin: a paradigm for the co-evolution of invariant TCRs and restricting MHCI-like molecules? Immunogenetics. 2016;68(8):537–48. Epub 2016/07/10. doi: 10.1007/s00251-016-0927-9. PubMed PMID: 27393664.

6. Gutierrez-Preciado A, Torres AG, Merino E, Bonomi HR, Goldbaum FA, Garcia-Angulo VA. Extensive Identification of Bacterial Riboflavin Transporters and Their Distribution across Bacterial Species. PLoS One. 2015;10(5):e0126124. Epub 2015/05/06. doi: 10.1371/journal.pone.0126124. PubMed PMID: 25938806; PubMed Central PMCID: PMCPMC4418817.

7. Corbett AJ, Eckle SB, Birkinshaw RW, Liu L, Patel O, Mahony J, et al. T-cell activation by transitory neo-antigens derived from distinct microbial pathways. Nature. 2014;509(7500):361–5. Epub 2014/04/04. doi: 10.1038/nature13160. PubMed PMID: 24695216.

8. Kjer-Nielsen L, Patel O, Corbett AJ, Le Nours J, Meehan B, Liu L, et al. MR1 presents microbial vitamin B metabolites to MAIT cells. Nature. 2012;491(7426):717–23. Epub 2012/10/12. doi: 10.1038/nature11605. PubMed PMID: 23051753.

9. Bacher A, Eberhardt S, Fischer M, Kis K, Richter G. Biosynthesis of vitamin b2 (riboflavin). Ann Rev Nutr. 2000;20:153–67. Epub 2000/08/15. doi: 10.1146/annurev.nutr.20.1.153. PubMed PMID: 10940330.

10. Tastan C, Karhan E, Zhou W, Fleming E, Voigt AY, Yao X, et al. Tuning of human MAIT cell activation by commensal bacteria species and MR1-dependent T-cell presentation. Mucosal Immunol. 2018;11(6):1591–605. Epub 2018/08/18. doi: 10.1038/s41385-018-0072-x. PubMed PMID: 30115998; PubMed Central PMCID: PMCPMC6279574.

11. Dusseaux M, Martin E, Serriari N, Peguillet I, Premel V, Louis D, et al. Human MAIT cells are xenobiotic-resistant, tissue-targeted, CD161hi IL-17-secreting T cells. Blood. 2011;117(4):1250–9. Epub 2010/11/19. doi: 10.1182/blood-2010-08-303339. PubMed PMID: 21084709.

12. Gibbs A, Leeansyah E, Introini A, Paquin-Proulx D, Hasselrot K, Andersson E, et al. MAIT cells reside in the female genital mucosa and are biased towards IL-17 and IL-22 production in response to bacterial stimulation. Mucosal Immunol. 2017;10(1):35–45. doi: 10.1038/mi.2016.30. PubMed PMID: 27049062; PubMed Central PMCID: PMC5053908.

13. Leeansyah E, Ganesh A, Quigley MF, Sonnerborg A, Andersson J, Hunt PW, et al. Activation, exhaustion, and persistent decline of the antimicrobial MR1-restricted MAIT-cell population in chronic HIV-1 infection. Blood. 2013;121(7):1124–35. doi: 10.1182/blood-2012-07-445429. PubMed PMID: 23243281; PubMed Central PMCID: PMC3575756.

14. Meierovics A, Yankelevich WJ, Cowley SC. MAIT cells are critical for optimal mucosal immune responses during in vivo pulmonary bacterial infection. Proc Natl Acad Sci USA. 2013;110(33):E3119–28. Epub 2013/07/31. doi: 10.1073/pnas.1302799110. PubMed PMID: 23898209; PubMed Central PMCID: PMCPMC3746930.

15. Constantinides MG, Link VM, Tamoutounour S, Wong AC, Perez-Chaparro PJ, Han SJ, et al. MAIT cells are imprinted by the microbiota in early life and promote tissue repair. Science. 2019;366(6464). Epub 2019/10/28. doi: 10.1126/science.aax6624. PubMed PMID: 31649166.

16. Hinks TSC, Marchi E, Jabeen M, Olshansky M, Kurioka A, Pediongco TJ, et al. Activation and In Vivo Evolution of the MAIT Cell Transcriptome in Mice and Humans Reveals Tissue Repair Functionality. Cell Rep. 2019;28(12):3249–62 e5. Epub 2019/09/19. doi: 10.1016/j.celrep.2019.07.039. PubMed PMID: 31533045.

17. Lamichhane R, Schneider M, de la Harpe SM, Harrop TWR, Hannaway RF, Dearden PK, et al. TCR- or Cytokine-Activated CD8(+) Mucosal-Associated Invariant T Cells Are Rapid Polyfunctional Effectors That Can Coordinate Immune Responses. Cell Rep. 2019;28(12):3061–76 e5. Epub 2019/09/19. doi: 10.1016/j.celrep.2019.08.054. PubMed PMID: 31533031.

18. Leng T, Akther HD, Hackstein CP, Powell K, King T, Friedrich M, et al. TCR and Inflammatory Signals Tune Human MAIT Cells to Exert Specific Tissue Repair and Effector Functions. Cell Rep. 2019;28(12):3077–91 e5. Epub 2019/09/19. doi: 10.1016/j.celrep.2019.08.050. PubMed PMID: 31533032.

19. Toubal A, Nel I, Lotersztajn S, Lehuen A. Mucosal-associated invariant T cells and disease. Nat Rev Immunol. 2019;19(10):643–57. Epub 2019/07/17. doi: 10.1038/s41577-019-0191-y. PubMed PMID: 31308521.

20. Nordmann P, Poirel L. Epidemiology and Diagnostics of Carbapenem Resistance in Gram-negative Bacteria. Clin Infect Dis. 2019;69(Supplement_7):S521–S8. Epub 2019/11/15. doi: 10.1093/cid/ciz824. PubMed PMID: 31724045; PubMed Central PMCID: PMCPMC6853758.

21. Macesic N, Nelson B, McConville TH, Giddins MJ, Green DA, Stump S, et al. Emergence of Polymyxin Resistance in Clinical Klebsiella pneumoniae Through Diverse Genetic Adaptations: A Genomic, Retrospective Cohort Study. Clin Infect Dis. 2019. Epub 2019/09/13. doi: 10.1093/cid/ciz623. PubMed PMID: 31513705.

22. Organization WH. Global priority list of antibiotic-resistant bacteria to guide research, discovery, and development of new antibiotics. Available at: http://www.whoint/medicines/publications/global-priority-list-antibiotic-resistant-bacteria/en/ Accessed 6 December 2019. 2017.

23. Lim TP, Wang R, Poh GQ, Koh TH, Tan TY, Lee W, et al. Integrated pharmacokinetic-pharmacodynamic modeling to evaluate empiric carbapenem therapy in bloodstream infections. Infect Drug Resist. 2018;11:1591–6. Epub 2018/10/13. doi: 10.2147/IDR.S168191. PubMed PMID: 30310294; PubMed Central PMCID: PMCPMC6166767.

24. Dias J, Sobkowiak MJ, Sandberg JK, Leeansyah E. Human MAIT-cell responses to Escherichia coli: activation, cytokine production, proliferation, and cytotoxicity. J Leukoc Biol. 2016;100(1):233–40. doi: 10.1189/jlb.4TA0815-391RR. PubMed PMID: 27034405; PubMed Central PMCID: PMC4946616.

25. Leeansyah E, Svard J, Dias J, Buggert M, Nystrom J, Quigley MF, et al. Arming of MAIT Cell Cytolytic Antimicrobial Activity Is Induced by IL-7 and Defective in HIV-1 Infection. PLoS Pathog. 2015;11(8):e1005072. Epub 2015/08/22. doi: 10.1371/journal.ppat.1005072. PubMed PMID: 26295709.

26. Giesubel U, Dalken B, Mahmud H, Wels WS. Cell binding, internalization and cytotoxic activity of human granzyme B expressed in the yeast Pichia pastoris. Biochem J. 2006;394(Pt 3):563–73. Epub 2005/12/13. doi: 10.1042/BJ20050687. PubMed PMID: 16336214; PubMed Central PMCID: PMCPMC1383706.

27. Vrazo AC, Hontz AE, Figueira SK, Butler BL, Ferrell JM, Binkowski BF, et al. Live cell evaluation of granzyme delivery and death receptor signaling in tumor cells targeted by human natural killer cells. Blood. 2015;126(8):e1–e10. Epub 2015/07/01. doi: 10.1182/blood-2015-03-632273. PubMed PMID: 26124495; PubMed Central PMCID: PMCPMC4543232.

28. Dotiwala F, Sen Santara S, Binker-Cosen AA, Li B, Chandrasekaran S, Lieberman J. Granzyme B Disrupts Central Metabolism and Protein Synthesis in Bacteria to Promote an Immune Cell Death Program. Cell. 2017;171(5):1125–37 e11. Epub 2017/11/07. doi: 10.1016/j.cell.2017.10.004. PubMed PMID: 29107333; PubMed Central PMCID: PMCPMC5693722.

29. Walch M, Dotiwala F, Mulik S, Thiery J, Kirchhausen T, Clayberger C, et al. Cytotoxic cells kill intracellular bacteria through granulysin-mediated delivery of granzymes. Cell. 2014;157(6):1309–23. Epub 2014/06/07. doi: 10.1016/j.cell.2014.03.062. PubMed PMID: 24906149; PubMed Central PMCID: PMCPMC4090916.

30. Wei HM, Lin LC, Wang CF, Lee YJ, Chen YT, Liao YD. Antimicrobial Properties of an Immunomodulator - 15 kDa Human Granulysin. PLoS One. 2016;11(6):e0156321. Epub 2016/06/09. doi: 10.1371/journal.pone.0156321. PubMed PMID: 27276051; PubMed Central PMCID: PMCPMC4898823.

31. Cai Y, Lim TP, Teo JQ, Sasikala S, Chan EC, Hong YJ, et al. Evaluating Polymyxin B-Based Combinations against Carbapenem-Resistant Escherichia coli in Time-Kill Studies and in a Hollow-Fiber Infection Model. Antimicrob Agents Chemother. 2017;61(1). Epub 2016/11/01. doi: 10.1128/AAC.01509-16. PubMed PMID: 27795375; PubMed Central PMCID: PMCPMC5192126.

32. Chua WJ, Truscott SM, Eickhoff CS, Blazevic A, Hoft DF, Hansen TH. Polyclonal mucosa-associated invariant T cells have unique innate functions in bacterial infection. Infect Immun. 2012;80(9):3256–67. Epub 2012/07/11. doi: 10.1128/iai.00279-12. PubMed PMID: 22778103; PubMed Central PMCID: PMCPMC3418730.

33. Kurioka A, Ussher JE, Cosgrove C, Clough C, Fergusson JR, Smith K, et al. MAIT cells are licensed through granzyme exchange to kill bacterially sensitized targets. Mucosal Immunol. 2015;8(2):429–40. Epub 2014/10/02. doi: 10.1038/mi.2014.81. PubMed PMID: 25269706; PubMed Central PMCID: PMCPMC4288950.

34. Le Bourhis L, Dusseaux M, Bohineust A, Bessoles S, Martin E, Premel V, et al. MAIT cells detect and efficiently lyse bacterially-infected epithelial cells. PLoS Pathog. 2013;9(10):e1003681. Epub 2013/10/17. doi: 10.1371/journal.ppat.1003681. PubMed PMID: 24130485; PubMed Central PMCID: PMCPMC3795036.

35. Meierovics AI, Cowley SC. MAIT cells promote inflammatory monocyte differentiation into dendritic cells during pulmonary intracellular infection. J Exp Med. 2016;213(12):2793–809. doi: 10.1084/jem.20160637. PubMed PMID: 27799620; PubMed Central PMCID: PMC5110023.

36. Wang H, D’Souza C, Lim XY, Kostenko L, Pediongco TJ, Eckle SBG, et al. MAIT cells protect against pulmonary Legionella longbeachae infection. Nat Commun. 2018;9(1):3350. Epub 2018/08/24. doi: 10.1038/s41467-018-05202-8. PubMed PMID: 30135490; PubMed Central PMCID: PMCPMC6105587.

37. Wang H, Kjer-Nielsen L, Shi M, D’Souza C, Pediongco TJ, Cao H, et al. IL-23 costimulates antigen-specific MAIT cell activation and enables vaccination against bacterial infection. Sci Immunol. 2019;4(41). Epub 2019/11/17. doi: 10.1126/sciimmunol.aaw0402. PubMed PMID: 31732518.

38. Balin SJ, Pellegrini M, Klechevsky E, Won ST, Weiss DI, Choi AW, et al. Human antimicrobial cytotoxic T lymphocytes, defined by NK receptors and antimicrobial proteins, kill intracellular bacteria. Sci Immunol. 2018;3(26). Epub 2018/09/02. doi: 10.1126/sciimmunol.aat7668. PubMed PMID: 30171080; PubMed Central PMCID: PMCPMC6431239.

39. Kaiserman D, Bird CH, Sun J, Matthews A, Ung K, Whisstock JC, et al. The major human and mouse granzymes are structurally and functionally divergent. J Cell Biol. 2006;175(4):619–30. Epub 2006/11/23. doi: 10.1083/jcb.200606073. PubMed PMID: 17116752; PubMed Central PMCID: PMCPMC2064598.

40. Dotiwala F, Lieberman J. Granulysin: killer lymphocyte safeguard against microbes. Curr Opin Immunol. 2019;60:19–29. Epub 2019/05/22. doi: 10.1016/j.coi.2019.04.013. PubMed PMID: 31112765; PubMed Central PMCID: PMCPMC6800608.

41. Jongstra J, Schall TJ, Dyer BJ, Clayberger C, Jorgensen J, Davis MM, et al. The isolation and sequence of a novel gene from a human functional T cell line. J Exp Med 1987;165(3):601–14. Epub 1987/03/01. doi: 10.1084/jem.165.3.601. PubMed PMID: 2434598; PubMed Central PMCID: PMCPMC2188281.

42. Sandberg JK, Norrby-Teglund A, Leeansyah E. Bacterial deception of MAIT cells in a cloud of superantigen and cytokines. PLoS Biol. 2017;15(7):e2003167. Epub 2017/07/26. doi: 10.1371/journal.pbio.2003167. PubMed PMID: 28742082; PubMed Central PMCID: PMCPMC5542701.

43. Shaler CR, Choi J, Rudak PT, Memarnejadian A, Szabo PA, Tun-Abraham ME, et al. MAIT cells launch a rapid, robust and distinct hyperinflammatory response to bacterial superantigens and quickly acquire an anergic phenotype that impedes their cognate antimicrobial function: Defining a novel mechanism of superantigen-induced immunopathology and immunosuppression. PLoS Biol. 2017;15(6):e2001930. doi: 10.1371/journal.pbio.2001930. PubMed PMID: 28632753; PubMed Central PMCID: PMC5478099.

44. Koay HF, Su S, Amann-Zalcenstein D, Daley SR, Comerford I, Miosge L, et al. A divergent transcriptional landscape underpins the development and functional branching of MAIT cells. Sci Immunol. 2019;4(41). Epub 2019/11/24. doi: 10.1126/sciimmunol.aay6039. PubMed PMID: 31757835.

45. Sobkowiak MJ, Davanian H, Heymann R, Gibbs A, Emgard J, Dias J, et al. Tissue-resident MAIT cell populations in human oral mucosa exhibit an activated profile and produce IL-17. Eur J Immunol. 2019;49(1):133–43. Epub 2018/10/30. doi: 10.1002/eji.201847759. PubMed PMID: 30372518.

46. Solders M, Gorchs L, Erkers T, Lundell AC, Nava S, Gidlof S, et al. MAIT cells accumulate in placental intervillous space and display a highly cytotoxic phenotype upon bacterial stimulation. Sci Rep. 2017;7(1):6123. Epub 2017/07/25. doi: 10.1038/s41598-017-06430-6. PubMed PMID: 28733576; PubMed Central PMCID: PMCPMC5522401.

47. Solders M, Gorchs L, Tiblad E, Gidlof S, Leeansyah E, Dias J, et al. Recruitment of MAIT Cells to the Intervillous Space of the Placenta by Placenta-Derived Chemokines. Front Immunol. 2019;10:1300. Epub 2019/06/28. doi: 10.3389/fimmu.2019.01300. PubMed PMID: 31244846; PubMed Central PMCID: PMCPMC6563723.

48. Ankomah P, Levin BR. Exploring the collaboration between antibiotics and the immune response in the treatment of acute, self-limiting infections. Proc Natl Acad Sci USA. 2014;111(23):8331–8. Epub 2014/05/21. doi: 10.1073/pnas.1400352111. PubMed PMID: 24843148; PubMed Central PMCID: PMCPMC4060691.

49. Gjini E, Brito PH. Integrating Antimicrobial Therapy with Host Immunity to Fight Drug-Resistant Infections: Classical vs. Adaptive Treatment. PLoS Comput Biol. 2016;12(4):e1004857. Epub 2016/04/15. doi: 10.1371/journal.pcbi.1004857. PubMed PMID: 27078624; PubMed Central PMCID: PMCPMC4831758.

50. Paupério FFS, Ganusov VV, Gjini E. Mathematical modeling links benefits of short and long antibiotic treatment to details of infection. bioRxiv. 2019:555334. doi: 10.1101/555334.

51. Handel A, Margolis E, Levin BR. Exploring the role of the immune response in preventing antibiotic resistance. J Theor Biol. 2009;256(4):655–62. Epub 2008/12/06. doi: 10.1016/j.jtbi.2008.10.025. PubMed PMID: 19056402; PubMed Central PMCID: PMCPMC5814249.

52. Magalhaes I, Pingris K, Poitou C, Bessoles S, Venteclef N, Kiaf B, et al. Mucosal-associated invariant T cell alterations in obese and type 2 diabetic patients. J Clin Invest. 2015;125(4):1752–62. Epub 2015/03/10. doi: 10.1172/jci78941. PubMed PMID: 25751065; PubMed Central PMCID: PMCPMC4396481.

53. Riva A, Patel V, Kurioka A, Jeffery HC, Wright G, Tarff S, et al. Mucosa-associated invariant T cells link intestinal immunity with antibacterial immune defects in alcoholic liver disease. Gut. 2018;67(5):918–30. Epub 2017/11/04. doi: 10.1136/gutjnl-2017-314458. PubMed PMID: 29097439; PubMed Central PMCID: PMCPMC5890654.

54. Rouxel O, Da Silva J, Beaudoin L, Nel I, Tard C, Cagninacci L, et al. Cytotoxic and regulatory roles of mucosal-associated invariant T cells in type 1 diabetes. Nat Immunol. 2017;18(12):1321–31. Epub 2017/10/11. doi: 10.1038/ni.3854. PubMed PMID: 28991267; PubMed Central PMCID: PMCPMC6025738.

55. Touch S, Assmann KE, Aron-Wisnewsky J, Marquet F, Rouault C, Fradet M, et al. Mucosal-associated invariant T (MAIT) cells are depleted and prone to apoptosis in cardiometabolic disorders. FASEB J. 2018:fj201800052RR. Epub 2018/06/30. doi: 10.1096/fj.201800052RR. PubMed PMID: 29957059.

56. Laudisio A, Marinosci F, Gemma A, Bartoli IR, Montenegro N, Incalzi RA. The Burden of Comorbidity Is Associated with Antibiotic Resistance Among Institutionalized Elderly with Urinary Infection: A Retrospective Cohort Study in a Single Italian Nursing Home Between 2009 and 2014. Microb Drug Resist. 2017;23(4):500–6. Epub 2016/08/16. doi: 10.1089/mdr.2016.0016. PubMed PMID: 27525808.

57. Miller BM, Johnson SW. Demographic and infection characteristics of patients with carbapenem-resistant Enterobacteriaceae in a community hospital: Development of a bedside clinical score for risk assessment. Am J Infect Control. 2016;44(2):134–7. Epub 2015/10/24. doi: 10.1016/j.ajic.2015.09.006. PubMed PMID: 26492818.

58. Nouvenne A, Ticinesi A, Lauretani F, Maggio M, Lippi G, Guida L, et al. Comorbidities and disease severity as risk factors for carbapenem-resistant Klebsiella pneumoniae colonization: report of an experience in an internal medicine unit. PLoS One. 2014;9(10):e110001. Epub 2014/10/22. doi: 10.1371/journal.pone.0110001. PubMed PMID: 25335100; PubMed Central PMCID: PMCPMC4198186.

59. Pouch SM, Satlin MJ. Carbapenem-resistant Enterobacteriaceae in special populations: Solid organ transplant recipients, stem cell transplant recipients, and patients with hematologic malignancies. Virulence. 2017;8(4):391–402. Epub 2016/07/30. doi: 10.1080/21505594.2016.1213472. PubMed PMID: 27470662; PubMed Central PMCID: PMCPMC5477691.

60. Ernst WA, Thoma-Uszynski S, Teitelbaum R, Ko C, Hanson DA, Clayberger C, et al. Granulysin, a T cell product, kills bacteria by altering membrane permeability. J Immunol. 2000;165(12):7102–8. Epub 2000/12/20. doi: 10.4049/jimmunol.165.12.7102. PubMed PMID: 11120840.

61. Krensky AM, Clayberger C. Biology and clinical relevance of granulysin. Tissue Antigens. 2009;73(3):193–8. Epub 2009/03/04. doi: 10.1111/j.1399-0039.2008.01218.x. PubMed PMID: 19254247; PubMed Central PMCID: PMCPMC2679253.

62. Stenger S, Hanson DA, Teitelbaum R, Dewan P, Niazi KR, Froelich CJ, et al. An antimicrobial activity of cytolytic T cells mediated by granulysin. Science. 1998;282(5386):121–5. Epub 1998/10/02. doi: 10.1126/science.282.5386.121. PubMed PMID: 9756476.

63. Dias J, Sandberg JK, Leeansyah E. Extensive Phenotypic Analysis, Transcription Factor Profiling, and Effector Cytokine Production of Human MAIT Cells by Flow Cytometry. Methods Mol Biol. 2017;1514:241–56. doi: 10.1007/978-1-4939-6548-9_17. PubMed PMID: 27787804.

64. Sia WR, Boulouis C, Gulam MY, Kwa ALH, Sandberg JK, Leeansyah E. Quantification of Human MAIT Cell-Mediated Cellular Cytotoxicity and Antimicrobial Activity. Methods Mol Biol. 2020;2098:149–65. Epub 2019/12/04. doi: 10.1007/978-1-0716-0207-2_10. PubMed PMID: 31792821.

65. Teo JQ, Chang CW, Leck H, Tang CY, Lee SJ, Cai Y, et al. Risk factors and outcomes associated with the isolation of polymyxin B and carbapenem-resistant Enterobacteriaceae spp.: A case-control study. Int J Antimicrob Agents. 2019;53(5):657–62. Epub 2019/03/19. doi: 10.1016/j.ijantimicag.2019.03.011. PubMed PMID: 30880229.

66. Zankari E, Hasman H, Cosentino S, Vestergaard M, Rasmussen S, Lund O, et al. Identification of acquired antimicrobial resistance genes. J Antimicrob Chemother. 2012;67(11):2640–4. Epub 2012/07/12. doi: 10.1093/jac/dks261. PubMed PMID: 22782487; PubMed Central PMCID: PMCPMC3468078.

67. Davies TA, Marie Queenan A, Morrow BJ, Shang W, Amsler K, He W, et al. Longitudinal survey of carbapenem resistance and resistance mechanisms in Enterobacteriaceae and non-fermenters from the USA in 2007-09. J Antimicrob Chemother. 2011;66(10):2298–307. Epub 2011/07/22. doi: 10.1093/jac/dkr290. PubMed PMID: 21775338.

68. Mak JY, Xu W, Reid RC, Corbett AJ, Meehan BS, Wang H, et al. Stabilizing short-lived Schiff base derivatives of 5-aminouracils that activate mucosal-associated invariant T cells. Nat Commun. 2017;8:14599. Epub 2017/03/09. doi: 10.1038/ncomms14599. PubMed PMID: 28272391; PubMed Central PMCID: PMCPMC5344979.

69. Roederer M, Nozzi JL, Nason MC. SPICE: exploration and analysis of post-cytometric complex multivariate datasets. Cytometry A. 2011;79(2):167–74. Epub 2011/01/26. doi: 10.1002/cyto.a.21015. PubMed PMID: 21265010; PubMed Central PMCID: PMCPMC3072288.

